# The Personalized Cancer Network Explorer (PeCaX) as a visual analytics tool to support molecular tumor boards

**DOI:** 10.1101/2021.06.25.449889

**Authors:** Mirjam Figaschewski, Bilge Sürün, Thorsten Tiede, Oliver Kohlbacher

## Abstract

**Background:** Personalized oncology represents a shift in cancer treatment from conventional methods to target specific therapies where the decisions are made based on the patient specific tumor profile. Selection of the optimal therapy relies on a complex interdisciplinary analysis and interpretation of these variants by experts in molecular tumor boards. With up to hundreds of somatic variants identified in a tumor, this process requires visual analytics tools to guide and accelerate the annotation process.

**Results:** The Personal Cancer Network Explorer (PeCaX) is a visual analytics tool supporting the efficient annotation, navigation, and interpretation of somatic genomic variants through functional annotation, drug target annotation, and visual interpretation within the context of biological networks. Starting with somatic variants in a VCF file, PeCaX enables users to explore these variants through a web-based graphical user interface. The most protruding feature of PeCaX is the combination of clinical variant annotation and gene-drug networks with an interactive visualization. This reduces the time and effort the user needs to invest to get to a treatment suggestion and helps to generate new hypotheses. PeCaX is being provided as a platform-independent containerized software package for local or institution-wide deployment. PeCaX is available for download at https://github.com/KohlbacherLab/PeCaX-docker.

## Background

Cancer is typically caused by genomic alterations inducing unchecked cellular proliferation. In personalized oncology [6], molecular data (e.g., genomics) is used jointly with clinical data to stratify therapies and choose the therapy best-suited for a specific patient. Next Generation Sequencing (NGS) is widely used to find those genomic alterations, such as single-nucleotide variants (SNVs), copy number variations (CNVs), or gene fusions.

Based on this data, the typical analysis workflow is usually as follows:

1. The (cancer) genome of a patient is sequenced and the SNVs and CNVs are stored in a Variant Call Format (VCF) file.
2. Variants are annotated with their effect.
3. Usually only variants with a strong effect are considered.
4. The remaining variants are looked up in databases to identify driver genes and to find drugs associated with these potential targets.
5. If no drug can be found for this specific variant or it is not applicable for the patient, pathways containing the related gene are considered to find druggable targets up- or downstream of the actionable variant.

Numerous tools exist for displaying and storing the information of a VCF file in the common tab separated values format (step 1) and to filter the variants for given annotations (step 3), e.g. VCF-Miner [4], BrowseVCF [14], VCF-Explorer [1]. But only few applications include the analysis of the SNVs and CNVs and the annotation of the variant effect (step 2), e.g. VCF-Server [7]. Perera-Bel *et al.* offer the additional option to find drugs targeting the variants (step 4) but their method does not perform variant effect prediction (step 2) and is limited to a specific data structure [12]. It also lacks a graphical user interface (GUI). OncoPDSS performs steps 1 to 4 but it is a web-server which can be a data security issue. Therefore, OncoPDSS does not store the input or results of the analysis [19]. These are displayed in unsearchable tables focusing on information about available pharmacotherapies which can be downloaded as TSV files. But it does not give information on the pathway context of a gene. So far, this information has to be collected manually.

We present PeCaX (Personalized Cancer Network Explorer), an integrated application for personalized oncology workflows. PeCaX performs clinical variant annotation by processing SNVs and CNVs and identifying clinically relevant variants and their targeting therapeutics using ClinVAP [15]. Networks containing the connections between the driver genes and the genes in their neighborhood as well as drugs targeting genes in this network are created through SBML4j [17] and they are visualized with the use of BioGraphVisart [11]. Our user-friendly, web-based user interface does not only interactively display the report generated by ClinVAP and the networks, but adds web links to external gene and drug databases and gives the option to take notes which are stored in PeCaX along with the information presented in the tables and networks. This allows the user to interactively work on and present the results, e.g. in a Molecular Tumor Board (MTB). The report can be downloaded as PDFs. In addition, the networks are available for download in publication-ready file formats (PNG, SVG and GraphML). In contrast to many VCF-analysis and clinical decision support tools, PeCaX is a local application with a graphical user interface working on the user’s local machine avoiding data privacy issues arising from the use of cloud-based services.

## Implementation

### Technical overview

PeCaX is a service-oriented local application and has been built using the NuxtJS framework for its web-based front end and integrates several other local services developed by us via REST APIs (see Fig. 1). It was developed in close interaction with clinical users to ensure a user-friendly interface. It supports concurrent use and works on any modern web browser independent of the operating system. It is easy to deploy via pre-built docker containers and easily integrated using docker compose.

**Figure 1:**
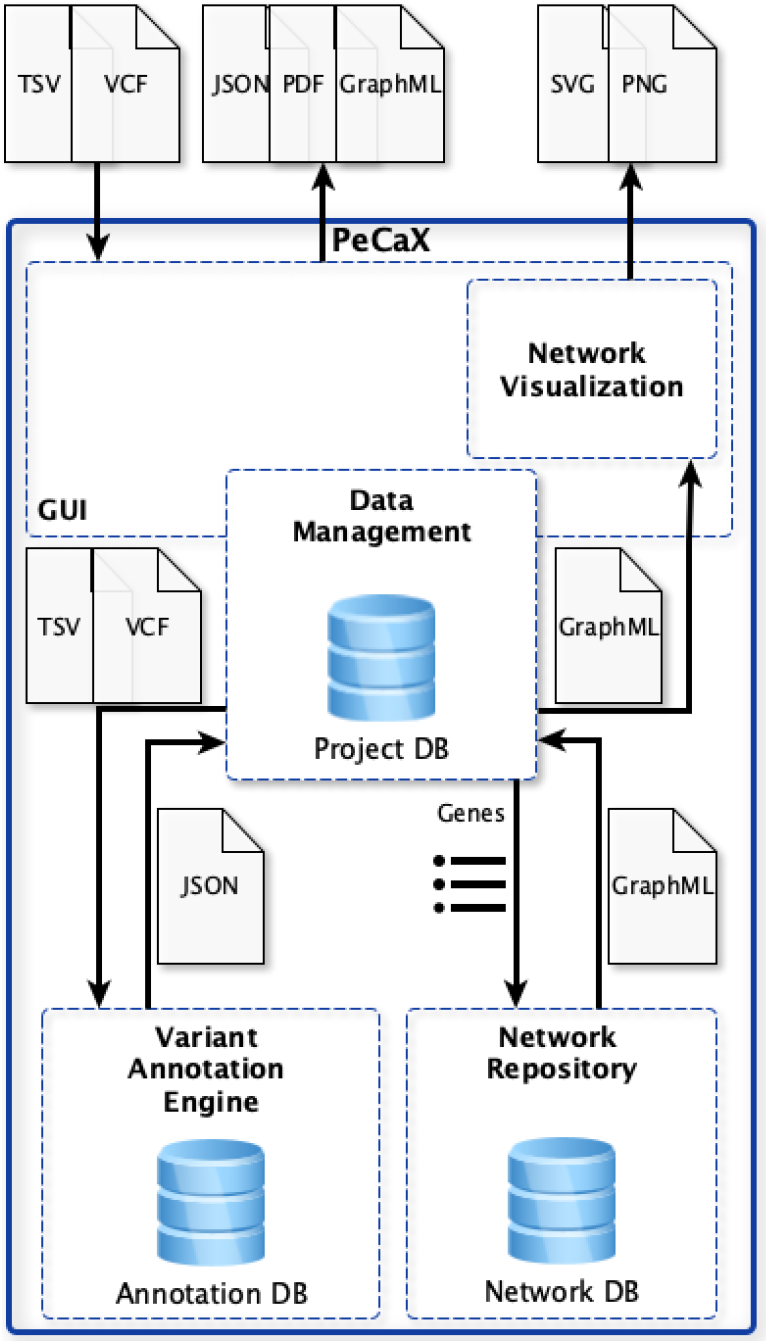
Architecture of PeCaX. Databases are indicated by the database icon.

### Clinical variant annotation

PeCaX relies on VCF files for information on SNVs and TSV files for optional CNV information. It integrates ClinVAP [15] to create a case report by processing variants using functional and clinical annotations of the genomic aberrations observed in a patient. ClinVAP employs Ensemble Variant Effect Predictor (VEP) [10] to obtain functional effects of the observed variants and filters them based on the severity of the predictions. It also performs clinical annotation to reveal driver genes and actionable targets using an integrated knowledge-base from publicly available databases (e.g., COSMIC [3], CGI [16]). As soon as the annotation is finished the variant files are deleted and PeCaX receives the resulting report as JSON file with information structured into five categories: known driver genes, drugs targeting the variants, therapeutics targeting the affected genes, cancer drugs targeting the mutated genes, and drugs with known adverse effects.

### Network generation

The analysis of gene networks and their known targeting drugs help to generate new hypotheses for the understanding and treatment of diseases. Thus, PeCaX sends the list of genes of each category to SBML4j [17] which is a service for persisting biological models and pathways in SBML format in a graph database. This graph database is used to extract information on the local network context of each candidate gene. Based on cancer-related pathways from KEGG [8], PeCaX can thus infer which related genes are up-and downstream of a candidate gene with respect to gene regulation and signalling. Additionally, SBML4j gives information about drugs associated with any of the candidate as well as up-/downstream genes.

### Interactive graphical user interface

PeCaX provides a simple graphical user interface to upload variants and display the results. The report generated by ClinVAP is displayed in an interactive tabular form next to the networks generated by SBML4j.

For an easier analysis of the networks, they are visualized as network graphs using BioGraphVisart [11]. The goal of the network analysis is to not only see the individual component but also the local neighborhood crosstalk with known pathways, and nearby options for therapeutic intervention (druggable genes). BioGraphVisart is a web-based tool written in Javascript. It automates the layout of the network graph, the labeling of nodes (genes, drugs) and edges (interactions), the edge style for different interaction types, the node coloring according to easily modifiable node attributes, and the generation of legends. In addition, human genes and proteins can be grouped with respect to predefined pathways from KEGG.

### Data Management

The results are stored in a local database (ArangoDB). For each uploaded VCF file a new database entry is created containing information about the user-selected project name, the job ID, parameters set for the clinical variant annotation, IDs of the networks generated and stored in SBML4j, and the information contained in the tables displayed on the front end.

## Results

### Overview of PeCaX

PeCaX is a comprehensive GUI-based clinical decision support tool that requires no programming knowledge. Users can perform clinical annotation and gene drug interaction network analysis via the interactive graphical interface. PeCaX provides data security as it comes in Docker containers and all analysis are performed on local infrastructure. It is supported by all modern web browsers across platforms. Hence, it is easily integrated into diagnostic and MTB workflows to investigate the relevance of single variants, complete cases or cohorts, e.g. from GWAS.

We provide a web page with example data for demonstration purpose only at https://pecax.informatik.uni-tuebingen.de.

### Data upload

To annotate and analyze a data set, PeCaX requires (local) submission of the data. All data is assigned to a specific *project*, a generic way of grouping data sets and results (e.g., one project per tumor board meeting or for one tumor entity). PeCaX requires one VCF file as input containing information on SNVs and (optionally) a TSV file containing information on CNVs. Details on format requirements and conventions are found in the online documentation of PeCaX. In addition the assembly of the human reference genome used in the mapping of the sequencing data is required (both GRCh37 and GRCh38 are supported). The clinical variant annotation can be filtered by a pre-selected cancer diagnosis given as ICD10 code. After uploading, the data is automatically submitted to a local instance of ClinVAP. In order to ensure data privacy, the VCF files are removed from the containers after processing by ClinVAP. The results of the variant annotation are stored in the project database in JSON format together with associated metadata (e.g., project name and the job ID). The user can also upload a previously downloaded JSON file which will skip ClinVAP or can enter the job ID of an analysis executed previously to directly get to the analysis part of the UI (see Figure 2). All job IDs of a given project are listed on a subpage where the user can select and delete them individually. Deletion of a job ID removes all information stored for this ID in the project database as well as the generated networks from the network database to ensure data privacy.

**Figure 2:**
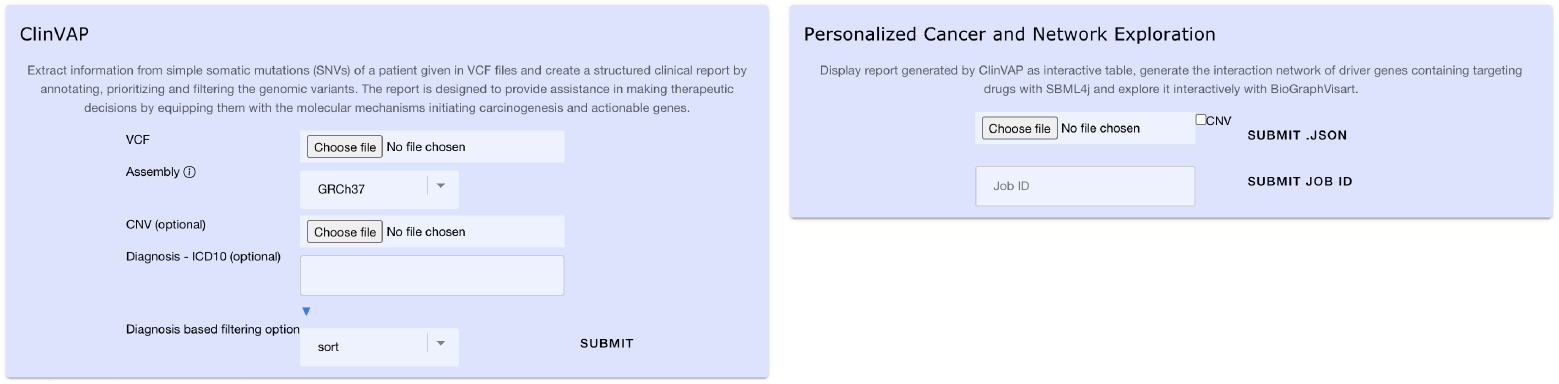
Screenshot of data upload.

### Interactive visualizations

The results of the clinical variant annotation performed is structured into several sections, which are all rendered as interactive, responsive tables. The first section contains the list of known cancer driver genes along with the somatic mutations observed in the patient. The list of drugs with the evidence of targeting a specific variant of the mutated gene and the documented drug response for the given mutational profile are displayed in the second section. The third section contains information on somatic mutations in pharmaceutical target affected genes and consists of two tables: therapies that have evidence of targeting the affected gene and the list of cancer drugs targeting the mutated gene. The fifth section contains the list of drugs with known adverse effects. References supporting the results found are displayed in a sixth section and all the somatic variants of the patient with their dbSNP and COSMIC IDs are listed in the last section.

Figure 3 shows a section of the results’ visualization. Each column of each table can be queried, filtered and sorted individually. The interactive table view supports a wide range of table operations in order to simplify navigation of the data, for example, highlighting of rows across sections, hiding/showing columns etc.

**Figure 3:**
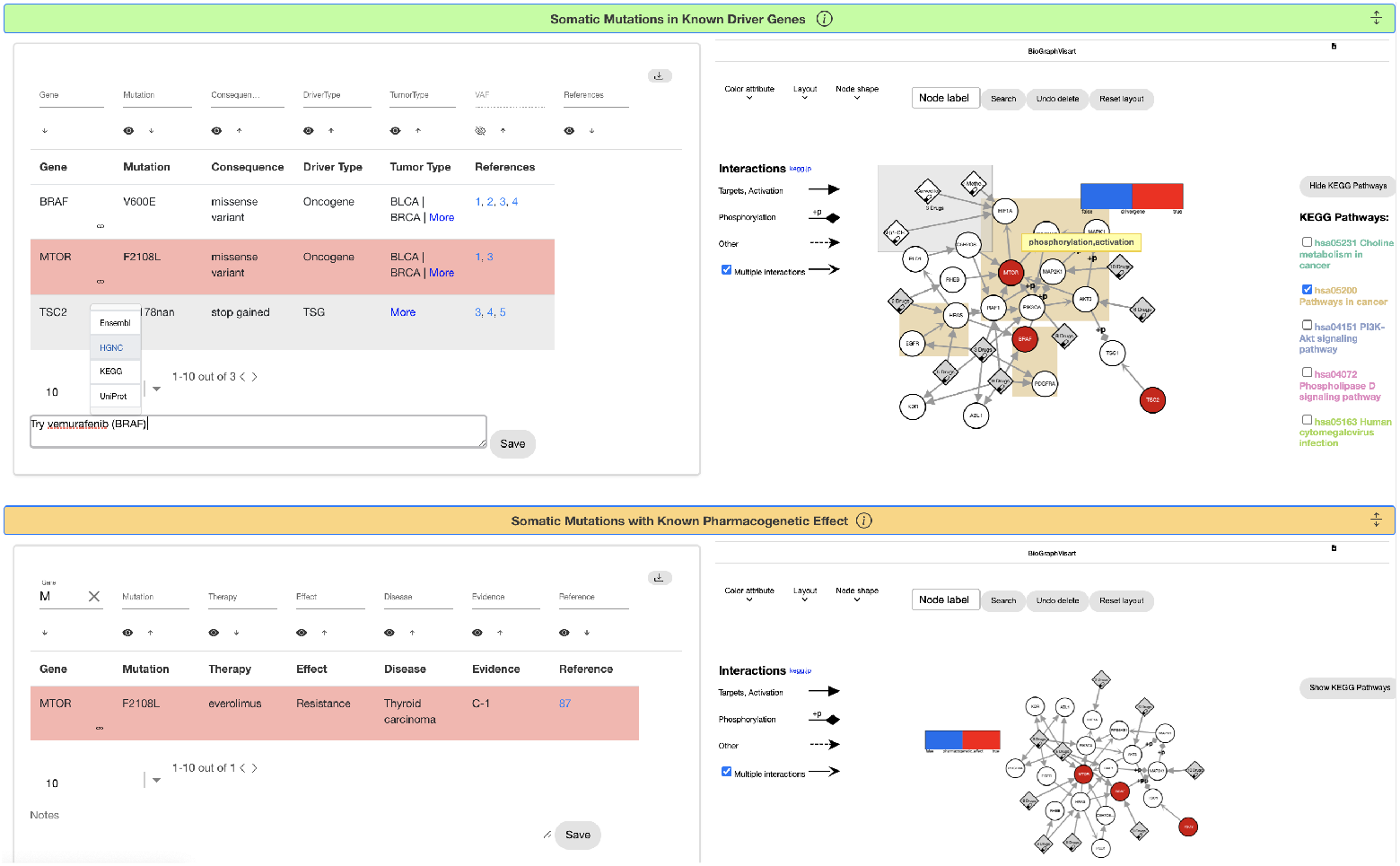
Screenshot of two tables (left) and related networks (right).

For each gene listed, the tables contain links to various external data sources such as Uniprot [2], KEGG [8] or Ensembl [5]. The links can be accessed via the drop-down menu next to the gene symbol and open in a separate browser tab or window. Likewise, the references given in the tables are directly linked to the web page of the related publication on PubMed or clinicaltrials.gov.

At the end of each section, users can add notes to be stored along with the annotated data in the internal database. These notes can be used to record conclusions from the analysis of the data and can be downloaded together with the table as PDF.

When at least one gene symbol of a table can be associated with an entry in the SBML4j database, a network for this table will be generated and is displayed next to it (see Fig. 3). The networks consist of nodes (genes, drugs) and edges (interactions). Genes found in the table are colored red and labelled with the gene symbol. Drugs associated with any of the genes in the network are represented as diamonds and are labelled with the drug name. If multiple drugs have the same gene target, they are merged into one node which is expandable in order to make the network representation visually more concise. Different interaction types (e.g., signaling, regulation) are depicted by different edge styles. If two nodes have multiple interactions, their edges are merged into one, which can be deactivated by the user. Since drug and gene names may become very long, they are shortened and moving the mouse over a node reveals the full node name. Edge types are treated in the same manner. The layout can be changed by the user based on five different layout types or the user can drag the nodes or the whole network manually to arrange them in the most informative layout. The nodes are searchable by the node label and deletable including connected edges. Gene nodes can be grouped and highlighted by associated KEGG pathways. A mouse click on a drug node links to an overview page with links to external databases containing information on this drug such as Drugbank [18], HGNC [13], and PDB [9].

### Exporting tables and networks

The clinical variant annotation report including the stored user-created notes and manual annotations can be downloaded as a whole in PDF and JSON format or every table individually in PDF format. The gene drug interaction networks are available for download individually in the formats PNG, SVG and GraphML.

## Conclusion

The individual nature of the genomic alterations causing cancer directly implies a personalized, or at least stratified approach to treating cancer if the underlying alterations are known. With the rapid drop in sequencing cost, sequencing has become routine for most cancers, but the interpretation of this data is still a major hurdle to clinical implementation of personalized oncology.

With PeCaX we present a novel tool for the exploration of the mutational landscape of a cancer patient and for treatment hypothesis generation. It is deployed in Docker containers guaranteeing full reproducibility independent of the operating system and as it is a local application it ensures data security and privacy. A local results database is used to keep track of the results and notes taken by the user. The combination of clinical variant annotation, gene drug interaction networks visualizing somatic variants in their pathway context and interactive web-based visualizations makes PeCaX unique and ensures ease of use for all users without the requirement of programming experience. PeCaX supports the diagnostic workflow, e.g. in a Molecular Tumor Board, to reach transparent personalized therapeutic decision in a shorter amount of time.

In the future, we plan to allow the upload of multiple VCF files at once for an easier comparison between patients. PeCaX might also include information on the patient’s background provided by the user, e.g., gender, age, and previous therapies.

